# Meditation Depth Enhances the Functional Signal-to-Noise Ratio of the Brain

**DOI:** 10.64898/2026.06.30.735351

**Authors:** Mihir Nath, Nicco Reggente, Neil Bailey, Morten Kringelbach, Ruben Laukkonen

## Abstract

Across contemplative traditions, deeper states of meditation are described as states of heightened clarity, vividness, and stillness of mind, yet what this clarity corresponds to in the brain has remained difficult to specify. The functional signal-to-noise ratio (f-SNR) framework frames mental clarity as a measurable property of neural signals: the degree to which brain activity tracks the causes of sensory signals rather than task-irrelevant endogenous activity. It predicts that deepening meditation should raise f-SNR, expressing sensory events more faithfully in neural signals against ongoing background activity. We tested this prediction across different levels of meditative depth. Twenty-nine experienced Vipassana practitioners meditated while auditory tones were presented, periodically reporting their depth of meditation. f-SNR was quantified from event-related potentials (ERPs) in a fronto-central P3 window and from single-trial decodability of auditory tone-evoked activity against no-tone background EEG. High-depth states were associated with greater ERP signal-to-noise ratio, stronger single-trial signal consistency, and improved decodability of sound tones. These results suggest that meditative depth is expressed in the reproducibility and stimulus-background separability of sensory responses, consistent with deep meditation enhancing the brain’s functional signal-to-noise ratio by improving the clarity of sensory signals and reducing task-irrelevant endogenous processes.

## Introduction

Reports of greater clarity, vividness, and immediacy of present-moment experience are among the most common descriptions of progressively deeper meditation across contemplative traditions [1, 2, 3]. Despite the consistency of this phenomenology in the literature on meditation, mental clarity has resisted precise neural operationalisation [4, 3]. Although often discussed as a characteristic of meditative phenomenology, and plausibly related to the attentional, affective, and perceptual changes associated with meditation practice [4, 2], phenomenological clarity has typically been described from the first-person perspective, with relatively little characterisation of the neurophysiological signatures that might underlie a clear mind [4, 2, 3]. Similar descriptors themselves recur often in research on meditation, including lucidity, vividness, stillness, and a sense of registering experience with less distortion [1, 2, 3]. These descriptions all seem to relate to the concept of signal quality. They therefore invite an empirical question: does the experience of a clearer mind correspond to a measurable change in how cleanly the brain represents the sensory events it encounters?

A recent theoretical proposal develops this possibility in terms of the brain’s functional signal-to-noise ratio (f-SNR) [5]. The theory begins from the premise that the brain is continuously modeling the causes of its sensory input [6, 7]. The fidelity of this modeling, understood as the degree to which neural activity tracks relevant causes while remaining free from task-irrelevant endogenous fluctuations, can be expressed as a functional signal-to-noise ratio [5]. In this account, f-SNR is high when neural activity selectively represents contextually relevant information, and low when activity is dominated by internal variability that does not improve perceptual or cognitive inference [5]. Meditation is proposed to raise f-SNR through two complementary operations: amplifying relevant signal and reducing the noise [5].

This proposal builds naturally on predictive-processing accounts of meditation. Predictive processing characterises perception as an active inferential process in which the brain generates predictions about the causes of sensory input and updates them in light of incoming evidence [6]. The relative influence of prior predictions and sensory signals depends on the precision assigned to each. Meditation practice may gradually recalibrate this balance, reducing the influence of habitual top-down predictions relative to present-moment sensory evidence [7]. Conceptual elaboration, self-referential interpretation, and learned evaluative reactions can all shape perception through such predictions. Although these processes are often adaptive, they can also filter, distort, or overdetermine experience [7]. With long-term meditation practice, the influence of these predictions may loosen, allowing present-moment sensory evidence to be registered with less top-down interference [7]. On this view, meditative clarity may reflect a state in which neural activity is more strongly coupled to present-moment input while being less dominated by internally generated predictions [5, 7]. One possible implication of this account is that f-SNR may increase not only across long-term training, but also as meditative states deepen within a single session of practice [5, 7].

The f-SNR framework already has a substantial empirical base, with several existing lines of meditation research converging on its central predictions [5]. Meditation experience has been associated with greater temporal stability of resting EEG dynamics [8], and intensive training has been shown to reduce reaction-time variability while increasing the consistency of evoked neural responses during an attention task [9]. Other work indicates that meditation can make the allocation of limited processing resources more efficient, including a reduced P3b to the first of two rapid targets alongside a smaller attentional blink [10], improved N2 and P3 markers of attentional control during a Stroop task [11], and improved perceptual discrimination during sustained attention [12]. Multivariate studies likewise suggest that meditation-related states have discriminable neural signatures. Internal attentional states can be decoded during meditation, with greater generalisability in experienced meditators [13], and distinct meditation styles can be classified in experts but not novices [14]. Viewed together, these otherwise separate findings cohere under a single construct, lending the f-SNR account convergent empirical support and motivating the more direct test pursued here.

If deeper meditation increases f-SNR, then even a simple, passive sensory event, requiring no decision or behavioural response, should be expressed more cleanly in neural activity as meditative depth increases [5]. Because such a stimulus does not require strategic task engagement, enhanced fidelity of its evoked response would point especially toward reduced endogenous interference: the decluttering of noise [5]. This prediction differs from much of the existing event-related literature on meditation, which has primarily examined how practitioners process events of differing salience or task relevance [4, 15, 16]. Experienced meditators have been reported to show attenuated responses to task-irrelevant distractors, consistent with reduced automatic orienting, and enhanced responses to targets when discrimination is required [17, 15, 16]. Such findings speak to how attention is allocated across salient, deviant, or goal-relevant events [15, 16]. They leave open a more fundamental question: independent of overt task demands, does the neural representation of sensory events become cleaner during meditation?

We addressed this question using data from experienced Vipassana practitioners meditating across two sessions while a passive auditory stream of stimuli was presented. The stream was a three-stimulus oddball paradigm adapted for meditation [15], termed the Sustained Auditory Passive Oddball (SAPO) task [18]: brief sounds occurred approximately once per second during extended periods of silent meditation, comprising a frequent standard tone, an infrequent pitch-deviant oddball, and an infrequent white-noise distractor. Our analysis focused on the standard tone, the most frequent and least demanding event in the stream, for two reasons. First, its high trial count affords the most stable estimate of the evoked response [19, 20]. Second, because it is expected, frequent, and response-irrelevant, it provides a conservative test of whether meditative depth improves the fidelity of passive sensory processing [15, 18].

Though conceptually distinct from recording-device SNR, f-SNR as a property of neural activity is nonetheless empirically related to it [19, 21, 5]. The more cleanly and stably the brain represents an event, the more recoverable that representation should be in a measurement system such as scalp EEG [19, 21, 5]. This connection makes standard event-related measures suitable proxies for f-SNR [19, 21, 20, 13]. We therefore quantified sensory signal fidelity using a convergent set of measures derived from standard-tone responses: ERP signal-to-noise ratio [21], single-trial consistency and split-half reliability of the ERP waveform [20], and decodability of tone-evoked activity from no-tone background EEG [13]. Together, these metrics capture different but related features of sensory signal fidelity, reflecting how clearly, consistently, and distinctly stimulus-evoked neural activity emerges from ongoing background activity [21, 20, 13].

Vipassana is a particularly suitable practice for testing the fSNR theory of meditation [1, 2, 5]. Vipassana practice trains sustained, non-reactive observation of present-moment sensory and somatic experience, cultivating equanimity toward whatever arises in awareness, rather than elaborating, suppressing, or identifying with it [1, 2]. Deepening Vipassana should therefore correspond to progressively less endogenous interference, creating the conditions under which the neural fidelity of sensory responses is expected to increase [5, 7]. Experienced practitioners in this tradition are also well placed to report moment-to-moment changes in the depth of their practice, given the central role of fine-grained introspective monitoring in Vippassana training [1].

Much of the meditation literature has relied on comparisons that treat meditation as a single, undifferentiated state, contrasting practitioners with non-practitioners or meditation with rest [4, 2]. Such designs have been productive, but they sit awkwardly alongside practitioners’ descriptions of meditation as a dynamic process that varies substantially within a single sitting [22, 23]. A practitioner may move through phases of clarity, dullness, stability, distraction, effort, and ease, rather than occupying one constant meditative state [22, 23]. The present dataset includes repeated self-reports of depth collected throughout extended meditation periods in the same individuals [18]. This allows f-SNR proxies to be related to meditative depth at the scale of moment-to-moment fluctuation, rather than only at the level of broad state or trait differences [18].

## Methods

### 2.1 Participants

The dataset analysed in the current work was drawn from the experiment originally described in Reggente et al. [18]. Participants were required to have a minimum of five years of continuous Vipassana meditation experience, with individual meditation sessions lasting no fewer than ten minutes and occurring at least five times per week. All participants attested to proficiency in the Vipassana technique as transmitted in the tradition of S.N. Goenka, which integrates breath-focused attentional training (*anāpānasati* ) with a systematic interoceptive body-scanning practice [1]. In this practice, awareness is iteratively directed through bodily regions in ascending and descending sequences, from the feet to the crown of the head and back again, while sustaining an equanimous, non-reactive stance toward all arising sensations.

Within the dataset, 37 participants completed data collection. Following EEG preprocessing and quality screening, 8 participants were excluded due to insufficient signal quality (details reported in section 2.3), yielding a final analytic sample of 29. This sample comprised 9 males and 20 females, ranging in age from 27 to 68 years (*M* = 46.28, *SD* = 12.69). On average, they had accumulated 15.52 years of meditation experience (*SD* = 11.62), maintained an average daily meditation frequency of 6.55 days per week (*SD* = 0.74), and had completed a cumulative total of 91.86 retreat days (*SD* = 191.48). Participants characterised their typical session length as follows: between 10 and 30 minutes (*n* = 6), between 30 minutes and one hour (*n* = 13), between one and two hours (*n* = 9), and more than two hours (*n* = 1).

### 2.2 Procedure

Before attending the laboratory session, participants received pre-study materials describing the experimental tasks. Upon arrival, they completed an initial questionnaire battery before physiological recording equipment was fitted. EEG was acquired using a 64-channel dual-amplifier BrainVision system (BrainVision LLC) sampled at 500 Hz. Head circumference was measured to select the appropriate cap size, and the cap was positioned with Cz at the vertex, verified by FPz at 10% of the nasion–inion distance. Skin at peripheral electrode sites (Skintact) was cleaned with isopropyl alcohol; for EEG cap electrodes, the scalp was lightly abraded with NuPrep gel before a high-chloride abrasive electrolyte gel (Neurospec; EasyCap, Inc.) was applied. Electrode impedances were monitored in BrainVision Recorder and maintained below 10 kΩ.

Peripheral physiological signals were recorded concurrently using the CGX-Aim II system (Cognionics Inc.), including electromyography (EMG), respiration, electrocardiography (ECG), and electrodermal activity (EDA). All data streams were synchronised online using NeuroPype Experiment Recorder (Intheon, La Jolla, CA) via the Lab Streaming Layer (LSL), which managed protocol timing and aligned EEG, physiological, and behavioural event streams within a common time base before saving them in Extensible Data Format (XDF).

During equipment setup, participants reviewed the task instructions, confirmed their understanding of meditation “depth” as a subjective construct, familiarised themselves with the clicker response device (used to indicate meditation depth, and confidence, explained later), and adjusted the audio volume to a comfortable level. They then moved to their assigned meditation space. To minimise power-line interference, all electronic devices were disconnected from wall power before recording commenced. Following the experimental session, recording equipment was removed and participants completed a final set of questionnaires.

#### 2.2.1 Experimental Sessions

Each participant completed two experimental sessions separated by at least one week. Each session comprised six sequential blocks (Figure 1): a 5-minute eyes-closed resting-state block without meditation; a 5-minute silent warm-up meditation block; a 35-minute meditation block with the SAPO task and periodic depth ratings; a 5-minute active oddball task requiring button-press responses to oddball tones; a second 35-minute meditation block without SAPO stimuli; and a 5-minute cool-down meditation block. Eyes remained closed throughout all blocks. Questionnaires were administered after each 35-minute meditation block.

**Figure 1:**
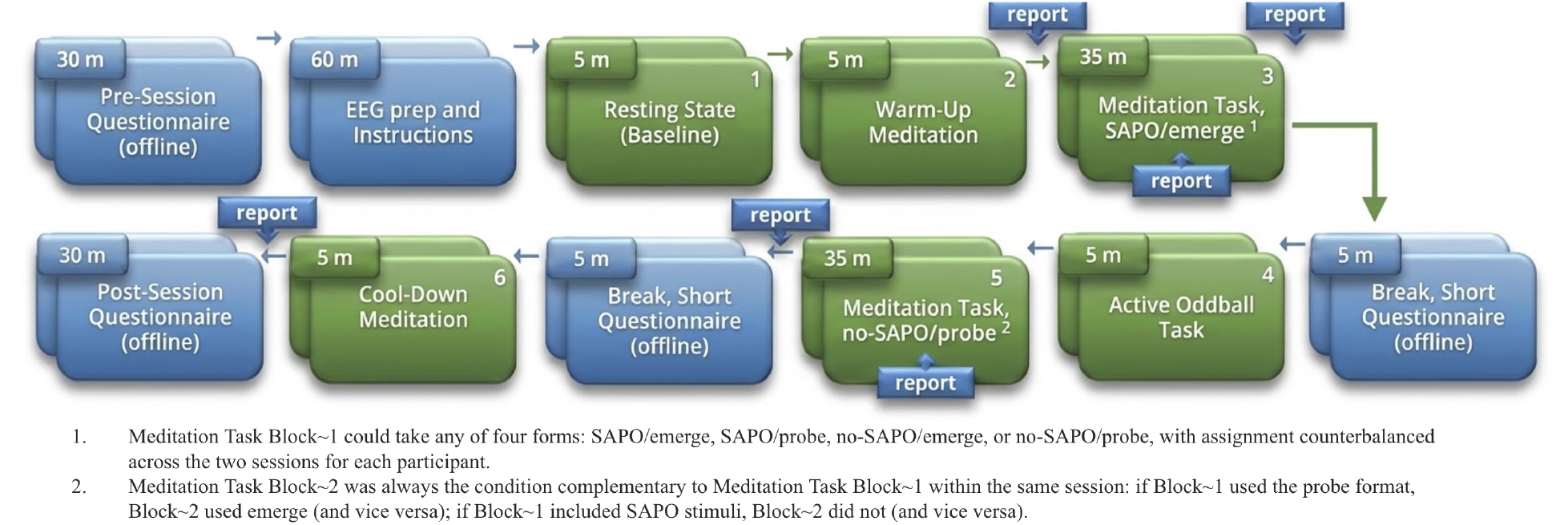
Experimental design for a representative session. Green blocks indicate segments during which EEG data were recorded; blue blocks denote segments completed offline without concurrent data acquisition. Downward arrows mark intervals at which depth reports were collected, while upward arrows indicate blocks in which continuous depth reporting occurred throughout.

**Figure 2:**
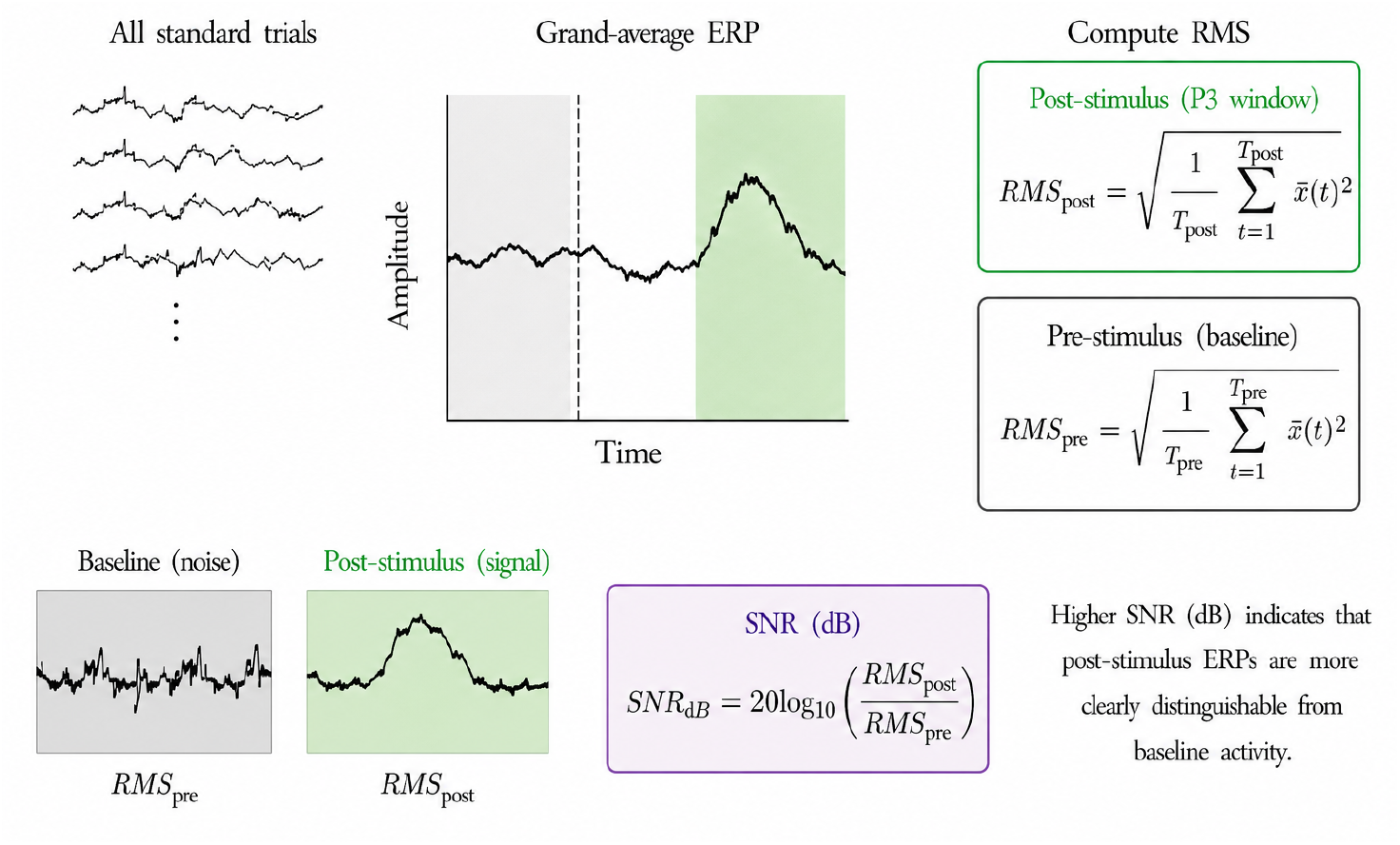
Schematic illustration of ERP signal-to-noise ratio computation.

The SAPO block was adapted from Cahn and colleagues’ passive three-stimulus auditory oddball meditation paradigm [15]. Participants continued meditating while task-irrelevant auditory stimuli were presented, comprising frequent standard tones, infrequent pitch-deviant oddballs, and infrequent white-noise distractors. Participants were instructed to maintain their meditation practice and make no behavioural responses, allowing stimulus-locked EEG responses to be recorded with minimal explicit task demands. The separate active oddball block used the same rare-target structure but required button-press responses to oddball tones, providing an active attentional comparison condition. The present analyses are restricted to the SAPO condition.

The two 35-minute meditation blocks used one of two depth-rating formats. In both, participants reported the greatest depth reached since the preceding report using a five-point clicker scale (1 = shallow or foundational; 5 = deepest or most culminative), followed by a confidence rating on the same scale. In the emerge format, participants submitted ratings spontaneously whenever they noticed that awareness had wandered, after which an auditory prompt elicited the confidence rating. In the probe format, depth and confidence ratings were elicited by auditory prompts occurring every 3 minutes to 3 minutes 50 seconds.

Across the two sessions, each participant completed all four combinations of stimulus condition (SAPO or no-SAPO) and rating format (probe or emerge): SAPO–probe, SAPO–emerge, no-SAPO– probe, and no-SAPO–emerge. Each combination was completed once, with the assignment of conditions to session and meditation block counterbalanced across participants.

### 2.3 Preprocessing

All preprocessing steps were implemented in Python 3.11 using MNE-Python, Autoreject, and SciPy [24, 25, 26]. The raw EEG signal was downsampled to 250 Hz and band-pass filtered between 0.5 and 45 Hz using a finite impulse response filter. Ocular artefacts, cardiac field interference, myogenic noise, and environmental disturbances were addressed using Independent Component Analysis (ICA) [27]. ICA was performed separately on each continuous recording session before segmentation, with artifactual components identified by visual inspection and projected out of the signal.

Following epoching, channel-level data quality was assessed using the Random Sample Consensus (RANSAC) algorithm implemented in Autoreject [28, 25]. RANSAC was configured with 50 resampling iterations, a minimum channel fraction of 0.25, a minimum reconstruction correlation of 0.85, and an unbroken-time criterion of 0.3. Segments containing more than 10 unreliable channels were excluded. In retained segments, channels identified as unreliable were marked as bad and spherically interpolated before further analysis.

Bad epochs were subsequently excluded using a custom multimetric rejection procedure based on root-mean-square (RMS) amplitude, peak-to-peak amplitude, and signal variance. RMS values were standardised across epochs within each segment. An epoch was rejected if its RMS exceeded two standard deviations above the segment mean, its peak-to-peak amplitude exceeded 150, or its signal variance fell below 5. Epochs meeting any one of these criteria were removed before further analysis.

### 2.4 Data Curation and Ground Truthing

The EEG signal was segmented by selecting a 60-second window preceding each depth report and assigning that window the participant’s five-point depth label. This duration balanced temporal specificity against the need for enough data to estimate event-related features reliably. For subsequent analyses, the five depth levels were mapped onto three broader depth categories: Low (levels 1 and 2), Moderate (level 3), and High (levels 4 and 5). This regrouping reduced sparsity across the original scale while preserving its ordinal structure and the distinction between shallow, intermediate, and deep meditative states. The endpoint of the selected window depended on the reporting condition. In the probe condition, the 60-second segment extended up to the time of the depth report itself. In the emerge condition, the end of the segment was instead set to 15 seconds before the depth report, so as to reduce the inclusion of neural activity associated with emergence from the reported state. Finally, only reports with confidence ratings at or above the midpoint of the five-point scale (i.e., a score of 3 or above) were retained.

### 2.5 Data Analysis

Functional signal-to-noise ratio (f-SNR) was operationalised with ERP measures of the clarity, stability, and consistency of auditory stimulus-locked responses [19, 29, 21]. We quantified f-SNR with four complementary metrics. Three were derived from the standard-tone ERP waveform: ERP signal-to-noise ratio (ERP-SNR), signal consistency, and split-half reliability. The waveform-based metrics used standard-tone epochs, which were the most numerous and therefore provided the most stable estimates [19]. Analyses focused on a fronto-central region of interest (Fz, F1, F2, F3, F4, FCz, FC1, FC2, FC3, FC4, Cz, C1, C2, C3, and C4) in the P3 window from 250 to 600 ms post-stimulus [29].

The fourth metric was classification-based: the decodability of the standard-tone response from no-tone background EEG, which indexes how distinctly the brain’s response to an ordinary tone could be separated from ongoing activity. As a secondary analysis, we additionally decoded between the auditory stimulus types themselves, contrasting standard, oddball, and distractor responses, to ask whether the ability to differentiate event types changed as a function of meditative depth.

#### 2.5.1 ERP Signal-to-Noise Ratio

ERP signal-to-noise ratio (ERP-SNR) was computed as the decibel-scaled ratio between the RMS amplitude of the averaged ERP in the P3 window and the RMS amplitude of the averaged prestimulus baseline [21]. Specifically,

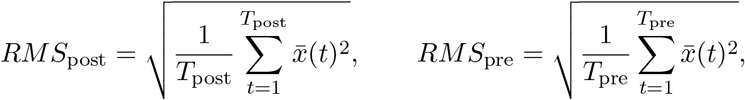

where *RMS*_post_ was computed over the analysis window and *RMS*_pre_ over the prestimulus baseline interval (*−*200 to 0 ms). SNR in decibels was then defined as

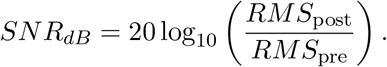

Higher values indicate greater separation between post-stimulus ERP activity and prestimulus baseline activity. During preprocessing, event-related epochs were retained without prestimulus baseline correction. Because prestimulus activity served as the noise reference in the ERP-SNR calculation, omitting baseline correction avoided coupling the pre-and post-stimulus intervals through subtraction of the same baseline mean.

#### 2.5.2 Signal Consistency

Signal consistency was computed as the mean Pearson correlation between each single trial and the averaged segment-level standard-tone ERP waveform from the same segment within the P3 analysis window [20]. For each trial,

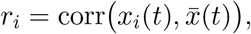

and signal consistency was defined as

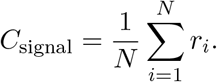

Because correlation is insensitive to absolute amplitude scaling, this measure primarily indexes morphological similarity between single trials and the grand-average response. Higher values therefore indicate that individual trials share a more consistent temporal structure.

#### 2.5.3 Split-Half Reliability

Split-half reliability quantified the internal reproducibility of the standard-tone ERP waveform within each meditation segment [30, 31, 20]. The standard-tone trials from each segment were divided into alternating odd-and even-indexed subsets, and an average ERP waveform was computed separately for each subset:

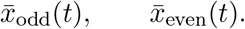

The Pearson correlation between the two half-average waveforms was then calculated:

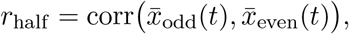

and adjusted using the Spearman–Brown prophecy formula [30, 31]:

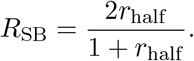

Higher values indicate that the segment-level ERP waveform was more reliably reproduced across independent subsets of standard-tone trials.

#### 2.5.4 Decodability

Decodability provided a complementary, classification-based index of f-SNR by quantifying how distinctly stimulus-evoked auditory activity could be separated from other EEG epochs. The primary f-SNR contrast distinguished standard-tone epochs, which were time-locked to auditory tone onset, from 1-second no-tone epochs randomly sampled from the same SAPO meditation blocks. The no-tone epochs were therefore drawn from the same experimental context and meditative-depth periods as the standard-tone epochs but were not time-locked to auditory events. This contrast indexed the separability of the stimulus-locked standard-tone response from ongoing EEG activity recorded during the same meditative condition.

As a secondary analysis, we decoded between the auditory stimulus types themselves using three event-type contrasts: standard versus oddball, standard versus distractor, and oddball versus distractor. These analyses tested whether the discriminability of different auditory events varied with meditative depth, rather than reflecting only the separation of stimulus-locked activity from ongoing EEG. All four contrasts used the same classification pipeline.

Classification was conducted separately for the Low, Moderate, and High depth groups using leave-one-subject-out cross-validation. Within each fold, the classifier was trained on data from all but one participant and evaluated on all available epochs from the held-out participant. Class balance was enforced in the training data by randomly undersampling the larger class to match the number of epochs in the smaller class.

Each epoch was independently reshaped from a channel-by-time matrix into a one-dimensional feature vector, with each feature corresponding to the EEG amplitude at a specific channel and time point. The resulting epoch-level vectors were stacked to form a two-dimensional matrix of epochs by channel–time features, which served as the input to the classification pipeline. This transformation preserved all channel and temporal samples and did not involve averaging across channels or time.

Within each fold, the mean and standard deviation of each channel–time feature were estimated exclusively from the training epochs. These training-derived parameters were used to mean-centre and scale both the training and held-out data, with no standardisation parameters estimated from the held-out participant. Principal component analysis (PCA) was subsequently fitted to the standardised training data and used to reduce the channel–time feature space to the top 50 orthogonal components [32]. Each component represented a weighted combination of the original channel–time features, yielding 50 component scores for each epoch. The held-out epochs were projected using the PCA loadings estimated from the training data only.

The resulting component scores were classified using a support vector machine with a radial basis function kernel [33]. The standardisation, PCA, and classifier were therefore fitted exclusively to the training data within each fold, preventing information from the held-out participant from influencing feature scaling, dimensionality reduction, or model fitting.

This procedure yielded one subject-level area under the receiver-operating characteristic curve (AUC) estimate for each held-out participant, depth group, and decoding contrast [34]. Higher AUC values indicate greater separability between the two epoch classes. Because oddball and distractor stimuli each comprised approximately 10% of trials, class balancing resulted in substantially fewer training epochs for the three event-type contrasts than for the standard-versus-no-tone contrast.

#### 2.5.5 Statistical Analysis

The waveform-based f-SNR metrics, namely ERP-SNR, signal consistency, and split-half reliability, were analysed using separate linear mixed-effects models with depth group specified as a categorical fixed effect and subject as a random intercept [35]. For each metric, an omnibus Wald chi-square test evaluated the main effect of depth group.

The four primary f-SNR measures comprised ERP-SNR, signal consistency, split-half reliability, and standard-versus-no-tone decodability. To control for multiple testing across these primary outcomes, FDR correction was applied jointly to the four corresponding omnibus *p*-values [36]. Where an omnibus effect was significant after this correction, the three pairwise contrasts between Low, Moderate, and High depth were obtained from the corresponding model, and their *p*-values were FDR-corrected as a separate family of three.

Thus, multiple-comparison correction was applied hierarchically at two levels: first across the four primary f-SNR omnibus tests, and subsequently across the three follow-up contrasts associated with each significant omnibus effect. Follow-up contrasts were corrected separately for each metric because they constituted conditional decompositions of that specific omnibus effect rather than independent research questions.

Each classification-based analysis used subject-level AUC values obtained from leave-one-subject-out cross-validation. For each decoding contrast, AUC values were clipped away from 0 and 1, logit-transformed, and entered into a linear mixed-effects model with depth group specified as a categorical fixed effect and subject as a random intercept. An omnibus Wald chi-square test evaluated the main effect of depth group. For the primary standard-versus-no-tone contrast, the omnibus *p*-value was included in the FDR correction across the four primary f-SNR measures described above. Where significant, the three pairwise contrasts between Low, Moderate, and High depth were obtained from the same model, with FDR correction applied to the resulting family of three *p*-values [36]. The same within-contrast correction procedure was applied to follow-up comparisons for the secondary decoding contrasts.

## 3 Results

We evaluated four complementary f-SNR measures: waveform-derived ERP-SNR, signal consistency, and split-half reliability, together with standard-versus-no-tone decodability. The waveform-derived metrics were computed from standard-tone ERPs in the frontocentral P3 window. These analyses included 572 standard-tone segments from 29 participants, comprising 229 Low-depth, 189 Moderate-depth, and 154 High-depth segment-level observations.

ERP-SNR differed significantly across depth groups (Figure 5a; *χ*^2^(2) = 13.89, *p*_FDR_ = .002). Mean ERP-SNR was comparable in Low depth (*M* = 5.58 dB) and Moderate depth (*M* = 5.54 dB), but was higher in High depth (*M* = 7.10 dB). FDR-corrected contrasts confirmed higher ERP-SNR in High than Low depth (*d* = 0.44, *p*_FDR_ = .001) and in High than Moderate depth (*d* = 0.34, *p*_FDR_ = .006).

**Figure 3:**
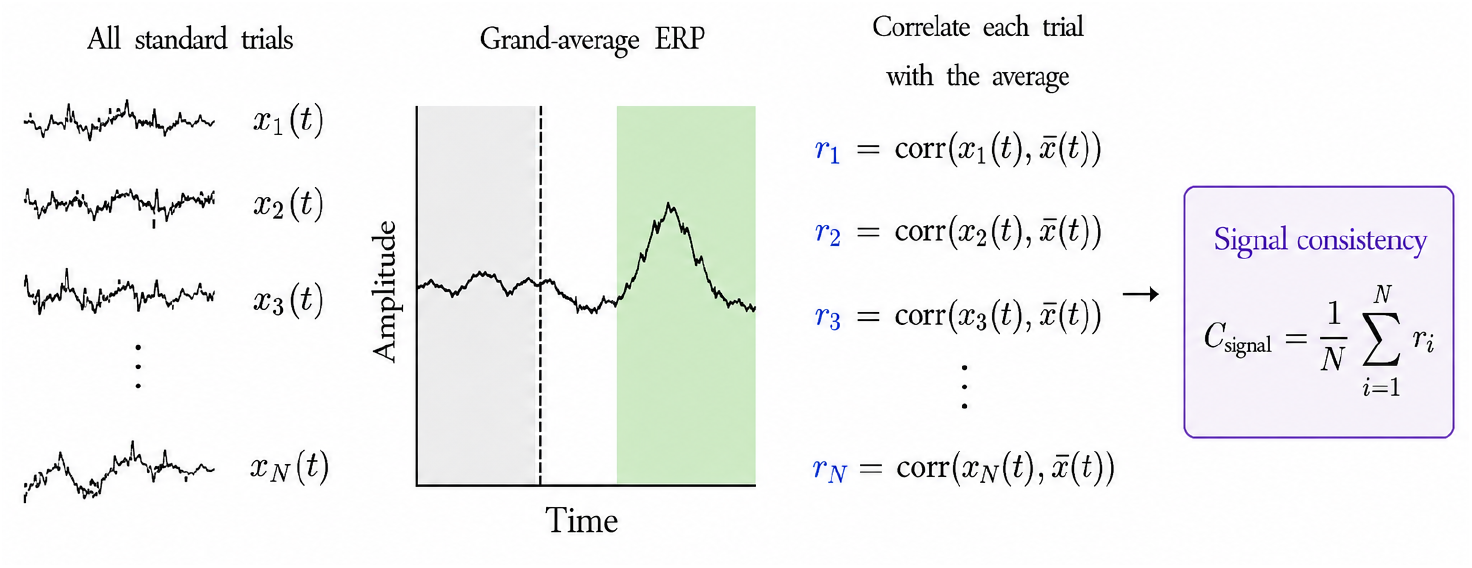
Schematic illustration of signal consistency computation.

**Figure 4:**
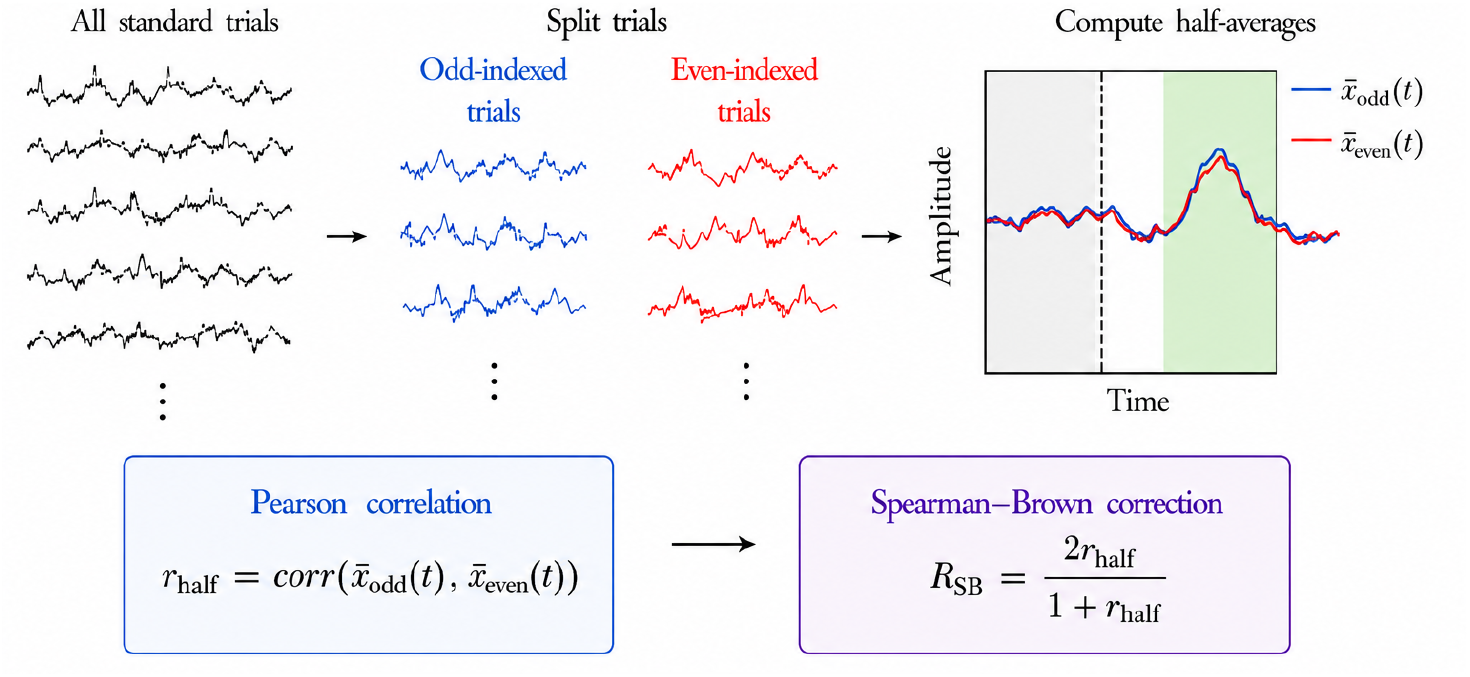
Schematic illustration of split-half reliability computation.

**Figure 5:**
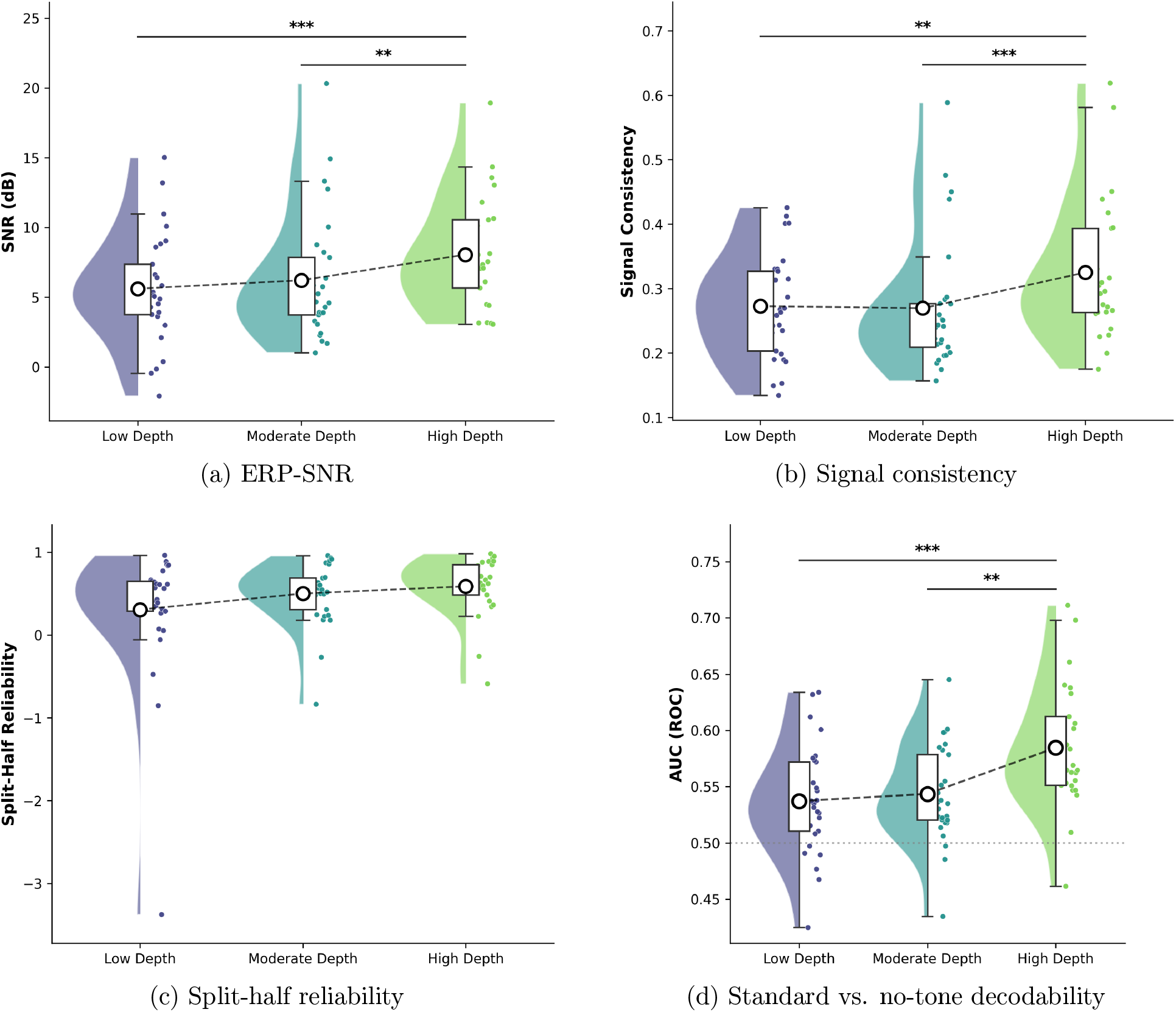
The four f-SNR measures across meditation depth. Panels show (a) ERP-SNR, (b) signal consistency, (c) split-half reliability, and (d) standard-versus-no-tone decodability. The first three are derived from the standard-tone ERP waveform; the fourth is the leave-one-subject-out decodability of the standard-tone response from no-tone background EEG.

Signal consistency showed the same depth-related pattern (Figure 5b; *χ*^2^(2) = 16.70, *p*_FDR_ *<* .001). Mean signal consistency was .273 in Low depth, .262 in Moderate depth, and .303 in High depth. High depth showed greater signal consistency than both Low depth (*d* = 0.35, *p*_FDR_ = .008) and Moderate depth (*d* = 0.47, *p*_FDR_ *<* .001).

Split-half reliability increased numerically across depth groups, from Low (*M* = .392) to Moderate (*M* = .455) and High (*M* = .495), but the omnibus depth effect did not reach the conventional significance threshold (Figure 5c; *χ*^2^(2) = 5.68, *p*_FDR_ = .058).

We then turned to the classification-based analyses, which indexed how distinctly stimulus-evoked activity could be separated from other epochs. We first tested whether standard-tone epochs could be distinguished from no-tone background epochs (Figure 5d). Before fold-specific undersampling, the depth-specific epoch pools contained 8,057 standard-tone and 10,752 background epochs in Low depth, 7,118 standard-tone and 9,012 background epochs in Moderate depth, and 5,925 standard-tone and 7,200 background epochs in High depth. Within each leave-one-subject-out fold, epochs from the held-out participant were first excluded from the training set, after which the larger remaining class was randomly undersampled to match the smaller class. The exact number of training epochs therefore varied across folds, but the two classes were balanced within every fold.

Mean AUC increased from Low (*M* = .543) and Moderate (*M* = .541) to High depth (*M* = .580). The omnibus mixed-effects model showed a significant effect of depth (*χ*^2^(2) = 9.83, *p*_FDR_ = .010). FDR-corrected contrasts showed greater decodability in High than Low depth (*d* = 1.02, *p*_FDR_ = .010) and in High than Moderate depth (*d* = 0.88, *p*_FDR_ = .006), whereas the Low–Moderate contrast was not significant (*d* = 0.14, *p*_FDR_ = .889).

As a secondary analysis, we asked whether meditation depth modulated decodability among the auditory stimulus types themselves, rather than between stimulus and background. The available epoch pools for the standard-versus-oddball contrast comprised 5,967 standard and 724 oddball epochs in Low depth, 5,216 standard and 660 oddball epochs in Moderate depth, and 4,356 standard and 610 oddball epochs in High depth. Because classification used balanced training sets, this contrast was capped at 610 epochs per class within each training fold. For standard-versus-distractor decoding, the corresponding pools contained 5,967/731, 5,216/654, and 4,356/598 standard/distractor epochs across Low, Moderate, and High depth, respectively, with training capped at 598 epochs per class. Oddball-versus-distractor decoding used 724/731, 660/654, and 610/598 oddball/distractor epochs, respectively, and was likewise capped at 598 epochs per class.

None of the three stimulus-type contrasts showed a significant depth effect after FDR correction across the three omnibus tests. Standard-versus-oddball decodability did not differ across depth (Low *M* = .656, Moderate *M* = .658, High *M* = .603; *χ*^2^(2) = 1.87, *p*_FDR_ = .392), nor did standard-versus-distractor decodability (Low *M* = .617, Moderate *M* = .636, High *M* = .578; *χ*^2^(2) = 3.51, *p*_FDR_ = .259) or oddball-versus-distractor decodability (Low *M* = .621, Moderate *M* = .509, High *M* = .539; *χ*^2^(2) = 6.19, *p*_FDR_ = .136).

Together, the results show that High-depth meditation was associated with a stronger and more internally consistent standard-tone response, alongside greater discriminability between standard-tone and no-tone epochs. The split-half reliability estimates followed the same numerical direction, but provided weaker evidence than ERP-SNR, signal consistency, and decodability. By contrast, decodability among the auditory stimulus types themselves did not vary reliably with depth, indicating that the depth-related effect was more evident for the separation of stimulus-evoked activity from ongoing background EEG than for discrimination among auditory event types.

## 4 Discussion

This study tested the general thesis that meditation makes the mind clearer by improving the signal-to-noise ratio of neural activations against the causes of sensory data. Each metric captured a different aspect of the stimulus evoked EEG response: amplitude-referenced clarity (ERP-SNR), morphological reproducibility (split-half reliability), trial-to-trial consistency (signal consistency), and separability from background activity (decodability). All four moved in the same direction as meditation depth increased. This convergence across complementary metrics is itself notable: it reduces the likelihood that any single result reflects a methodological artefact and instead supports a coherent underlying shift in the f-SNR of stimulus-evoked neural processing [5, 19, 20, 21].

Standard-versus-no-tone classification showed a robust depth effect, indicating that in High-depth meditation the ordinary standard-tone response was more separable from ongoing background EEG. By contrast, decodability among the auditory event types themselves did not vary reliably with depth. These null effects should be interpreted cautiously, because the oddball and distractor classes were relatively rare and class balancing left these secondary contrasts with substantially fewer training epochs than the standard-versus-no-tone analysis.

Two complementary mechanisms can increase f-SNR: amplification of contextually relevant signal and attenuation of the background activity that competes with it. Meditation may improve the fidelity of neural representations by simultaneously strengthening the representation of relevant sensory information and reducing interference from endogenous activity [5]. In the case of Vipassana, this corresponds theoretically to increasingly fine-grained tracking of sensory features alongside a reduction in conceptual overlay, reactive elaboration, and other internally generated activity that can obscure present-moment sensory evidence. The present findings are consistent with this dual-channel account.

A plausible neural substrate for this noise reduction is the suppression of default mode network (DMN) activity. The DMN is characterised by sustained activity during rest and mind-wandering and is reliably deactivated during externally directed or attentional tasks [37]. Critically, experienced meditators show reduced DMN activity during periods not only of rest, but also during other effortful cognitive tasks, suggesting that long-term practice may reduce self-referential and elaborative processing [38, 39]. From the perspective of the f-SNR theory, ongoing DMN-driven activity constitutes a source of endogenous neural interference that might compete with stimulus-driven responses in the EEG signal. If deeper meditative states suppress this source of variability more effectively, the stimulus-locked response would naturally emerge with greater clarity and reproducibility from the background, without any need to invoke enhanced sensory gain. This aligns with the proposal that deepening practice progressively targets deeper, increasingly self-related sources of endogenous noise, so that within-session increases in meditative depth should track reductions in precisely this kind of self-referential interference [5].

This account also aligns with the predictive-processing model of meditation, in which repeated practice is suggested to be associated with a withdrawal of precision from habitual top-down predictions, so that the dominance of higher-level predictive models over the processing of incoming evidence is progressively reduced [7]. As conceptual elaboration, evaluative reaction, and self-referential narrative exert less influence on lower-level sensory representations, stimulus-driven responses are expressed more faithfully and with less top-down distortion [7].

Beyond the theoretical implications, the f-SNR results explored here have significant translational potential. ERP-SNR can be computed from relatively short EEG segments and is therefore, in principle, amenable to near-real-time estimation. Signal consistency may similarly be estimated online if incoming trials are compared against a template established during an earlier calibration period or updated over the course of the session. Decodability is somewhat more challenging, as training a machine-learning classifier from scratch on short, continuously updating windows would require sufficient labelled data and introduce additional computational constraints. However, a classifier trained beforehand on baseline or calibration data could be held fixed and applied to short segments of incoming EEG, allowing stimulus-versus-background decodability to be estimated in near-real time. Together, these measures are therefore plausible candidates for closed-loop neurofeedback that could provide practitioners with continuous, objective information about their meditative state [40]. A validated real-time f-SNR index might guide instruction, flag mind-wandering before a practitioner notices it, or support a portable depth monitor for retreat, clinical, and training settings.

The same property connects to a further implication of the f-SNR account for brain–computer interfaces (BCIs). Much of BCI research has focused on improving the measurement and decoding side of the interface: developing better sensors, preprocessing methods, feature representations, and decoding algorithms to extract reliable information from inherently variable neural signals [41, 5]. The present findings point to a complementary possibility. If the decodability of neural activity depends partly on the functional state of the brain itself, then BCI performance might also be improved by increasing the stability and distinguishability of the neural signals being generated in the first place. In this view, meditation provides an intriguing proof of principle: deeper meditative states were associated with neural responses that were more consistent and more readily distinguishable from background activity. If such changes prove trainable and generalize to BCI-relevant signals, contemplative or other forms of mental training could potentially be used to place users in states that are intrinsically more amenable to neural decoding. This reframes part of the BCI problem from asking only how neural signals can be read more effectively to also asking whether users can learn to generate neural activity that is more reliably readable. Future work could therefore test whether experimentally increasing f-SNR improves decoder accuracy, calibration efficiency, or robustness across sessions, and whether such effects can be deliberately trained.

### 4.1 Limitations

Several limitations qualify these findings. Most fundamentally, meditative depth was defined by subjective self-report on a five-point scale, with no objective ground truth; although confidence filtering and within-participant modelling (provided by the linear mixed model) reduced the impact of idiosyncratic scale use, depth may not correspond to a single underlying neural dimension [22, 23]. A second concern is physiological: although ICA removed ocular, cardiac, and myogenic artefacts, residual differences in movement, muscle tone, or arousal across depth states could contribute to the apparent change in signal-to-noise, and the present data cannot fully separate recording that is less influenced by artifacts from a quieter neural background.

The study is also based on a modest sample of highly experienced practitioners from a single Vipassana tradition. It cannot establish that meditative depth as understood in other traditions would provide a similar result. Longitudinal, interventional, and cross-technique designs with larger and more balanced samples will be needed to establish the causal structure and boundary conditions of the effects reported here.

### 4.2 Conclusion

The current work examined whether subjective changes in meditative depth, as reported by experienced Vipassana practitioners, are accompanied by measurable changes in f-SNR. Across four complementary metrics, comprising ERP-SNR, single-trial signal consistency, split-half reliability, and standard-versus-no-tone decodability, deeper states were associated with consistently higher f-SNR. Deeper Vipassana appears to produce a processing regime in which ordinary auditory responses are more clearly and reproducibly expressed against ongoing neural activity. The consistency of the f-SNR effects and their coherence with predictive processing accounts of meditation provide a principled empirical foundation for the idea that a clear mind may be reflected in a clear brain. The present work offers both a methodological template and a set of candidate markers for future investigation into the neural signatures of meditation.

## Acknowledgements

We gratefully thank the study participants for their time and practice, and the researchers and interns at the Institute for Advanced Consciousness Studies who contributed to data collection.

